# Seed size control via phloem end by callose deposition/degradation of β-1,3-glucanase

**DOI:** 10.1101/2023.07.23.550179

**Authors:** Xiaoyan Liu, Kohdai P. Nakajima, Xiaoyan Wu, Shaowei Zhu, Prakash Babu Adhikari, Kentaro Okada, Ken-ichi Kurotani, Takashi Ishida, Masayoshi Nakamura, Yoshikatsu Sato, Liyang Xie, Chen Huang, Jiale He, Shinichiro Sawa, Tetsuya Higashiyama, Michitaka Notaguchi, Ryushiro D. Kasahara

## Abstract

Seed formation is crucial for lives of plants as well as humans; however, the mechanisms governing seed size require further investigation. Here, we present a new mechanism to modify the seed size by the newly identified phloem end that support nutrient transport, at the chalazal end of the ovule, however, blocked by callose deposition. Callose is removed after central cell fertilization (open state), allowing nutrients to be transported to the seed. However, if fertilization fails, callose deposition persists (closed state), preventing the phloem end from transporting nutrients. β-1,3-glucanase genes, including putative plasmodesmata-associated proteins (AtBG_ppap), were identified as regulators of callose removal. The *Atbg*_*ppap* mutant had the phloem end in the closed state and produced smaller seeds due to incomplete callose degradation. In contrast, the *AtBG_ppap* overexpression line produced larger seeds than the wild type due to continuous callose degradation, indicating that the phloem end regulates substance flow via callose deposition/degradation.

## Introduction

Seed size is a crucial trait for plant breeding. Although several genes required for normal seed size have been identified (Orozco-Arroyo et al., 2015; Li et al., 2019), the mechanisms underlying why and how most genes directly affect the phenotype are still unclear. Moreover, most of these genes are associated with either hormonal pathways, maternal control, or other broad regulatory responses, except for the amino acid transporter UmamiT (Müller et al., 2015). Plants with defective UmamiT produce smaller seeds than those with the wild type (WT), since the mutant cannot transport amino acids. Therefore, nutrition plays a critical role in seed size and offers huge potential for crop seed enlargement. In both animals and plants, nutrients supplied by the maternal bodies are essential for embryo formation and development. In plants, the transmitting tract (akin to the placenta in animals) and funiculus (akin to the umbilical cord in animals) are fully developed before fertilization, implying that plants have unique nutrition-support systems. However, the mechanism by which plants supply nutrients from the transmitting tract to developing seeds after fertilization is not well understood. While several studies have reported that fertilization is necessary for seed formation (Dumas and Rogowsky, 2008; Hands et al., 2016; Dresselhaus et al., 2016; Adhikari et al., 2020), with the exception of apomixis (Koltunow and Grossniklaus, 2003; Underwood and Mercier, 2022), none have been able to explain the need for fertilization to initiate or continue the nutrient transport required for normal seed formation. Similarly, studies that have identified nutrition transporters in the ovule (Stadler et al., 2005; Werner et al., 2011; Müller et al., 2015; Vogiatzaki et al., 2017; Karmann et al, 2018) were unable to explain how fertilization triggers ovule nutrition to induce seed formation. Generative Cell Specific *1 (gcs1*) mutant (Mori et al., 2006; von Besser et al., 2006; Nagahara et al., 2015), produces the pollen contained fertilization-defective sperm cells. The seed formation by the *gcs1* mutant, which was partially induced by the release of pollen tube contents into the ovule, was arrested during the pollen tube-dependent ovule enlargement morphology (POEM) (Kasahara et al., 2016; Kasahara et al., 2017). The POEM results indicated that a fertilization-dependent additional regulatory mechanism is required to complete seed formation.

In this study, we were able to show that (i) A new plant nutrition regulatory mechanism is governed by callose deposition/degradation during seed formation. Callose deposition increases if fertilization fails, but callose degrades if an ovule is successfully fertilized. (ii) The expression of the *AtBG_ppap* gene, encoding *β*-1,3-glucanase, is upregulated only after successful fertilization, and its expression is correlated with increased seed size. Defect in *AtBG_ppap* (*Atbg_ppap* mutant) leads to an increase in callose deposition, thereby interfering with nutrient supply to the developing seeds and negatively affecting their size. In contrast, overexpression of *AtBG_ppap* (*OEAtBG_ppap*) leads to an increase in seed size due to the smooth nutrition flow to the developing seed led by the increased callose degradation. (iii) Central cell fertilization is required for callose degradation and initiation of sucrose transport. (iv) The phloem-end structure was identified. So far, only branching structures have been reported in the early stage of the ovule before fertilization. Notably, the final form of the mature phloem was identified in this study.

## Results

### Discovery of the callose deposition phenomenon under failure of fertilization

Since callose accumulation in the leaf phloem reduces axillary bud growth (Paterlini et al., 2021), it could also significantly affect seed formation and development. In our study, we observed significant callose deposition around the chalazal end (cyan ellipses in Fig. 1) after aniline blue staining (Hough et al., 1985) at 2–4 days after emasculation (DAE) in no pollen tube (NPT) ovules (Fig. 1A–C). The signal was not observed at 2 DAE, which is equivalent to 1 d after pollination (DAP, Fig. 1A), but gradually increased at ≥3 DAE (Fig. 1B, C). In wild-type (WT) ovules, the signal was observed at 1 DAP (Fig. 1D) but became weaker at 2 DAP and 3 DAP (Fig. 1E, F), indicating that callose deposition was transiently promoted at 1 DAP (=2 DAE), which diminished later. In contrast, in the ovules crossed with *gcs1* mutant pollen (*gcs1* ovules) the signal remained consistently intense, even at ≥2 DAP (Fig. 1G–I). The signal intensities were further investigated both after fertilization (WT) and fertilization failure (*gcs1*) at every 4 hours (Fig. S1). At 24 hours after pollination (HAP) in WT, the signal intensity was starting decreased and at 48 HAP, the signal became weaker. In contrast, at 24 HAP in *gcs1* mutant, the signal became consistently intense until at 48 HAP. As mentioned earlier, the *gcs1* mutant does not produce seeds, because its sperm cells are defective and can fertilize neither egg nor central cells. Considering that the phenotypic difference in WT, against NPT and the *gcs1* mutant, depends on whether the ovule is fertilized or not, indicating that if the ovule completes fertilization, it does not accumulate callose at the chalazal end, but deposition continues if not fertilized. Even though the amount of callose deposition varied between NPT (Fig. 1A–C) and *gcs1* mutant (Fig. 1G–I) ovules, the level of callose deposition was conceptually identical; deposited callose was not removed if the ovule was not fertilized. To further investigate this phenomenon, three-dimensional images of the WT ovules (Fig. 1J–N, Video S1) and *gcs1* mutant ovules (Fig. 1O–S, Video S1) were captured at 2 DAP. In the WT ovules, the central region of the deposition was degraded, showing a ring-shaped signal, whereas *gcs1* mutant ovules displayed a saturated signal. However, both showed circular structures, suggesting that their basic structures were similar. These results indicate that callose deposition persists if ovule fertilization fails; in contrast, if it is fertilized, the deposits are degraded.

**Fig. 1.**
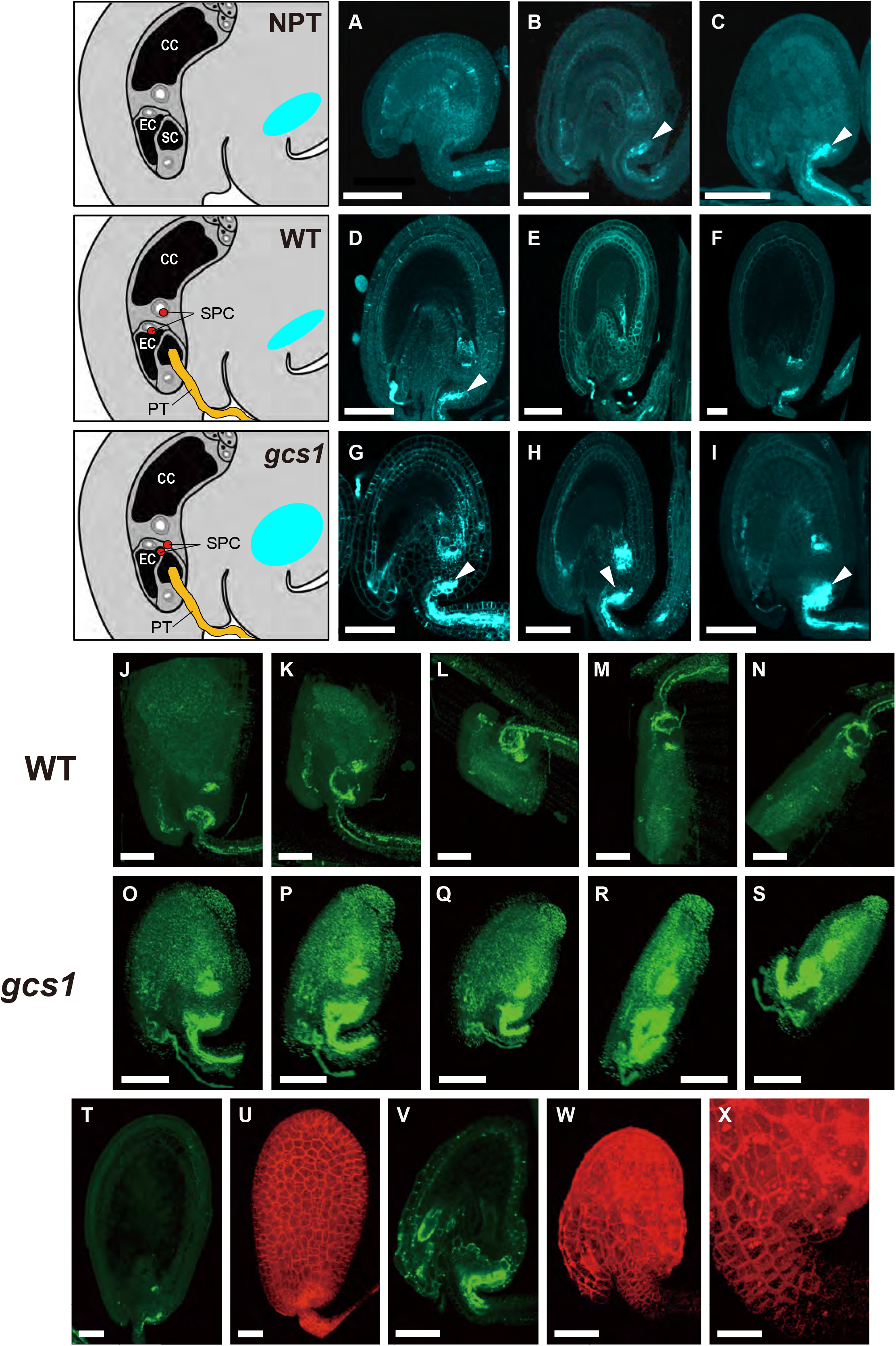
Discovery of the callose deposition phenomenon. Aniline blue staining of ovules with no pollen tube (NPT) at 2 DAE (**A**), 3 DAE (**B**), and 4 DAE (**C**). WT (wild-type background) ovules at 1 DAP (equivalent to 2 DAE) (**D**), 2 DAP (= 3 DAE) (**E**), and 3 DAP (= 4 DAE) (**F**). *gcs1* (*gcs1* background) mutant ovules at 1 DAP (**G**), 2 DAP (**H**), and 3 DAP (**I**). The cyan color indicates the position and intensity of the aniline blue-stained region. (**J**−**S**) Rotated images of callose deposition in the WT and *gcs1* mutant. (**J**–**N**) Aniline blue staining of WT ovules. Weak callose deposits were detected in a round shape at the chalazal end of the ovule. (**O**−**S**) Aniline blue staining of the *gcs1* mutant ovules. Intensive round callose depositions were identified at the chalazal end of the ovule. Aniline blue staining (**T**) and RPS5Apro::tdTomato-LTI6b expression (**U**) for WT at 3 DAP. Aniline blue staining (**V**) and RPS5Apro::tdTomato-LTI6b expression (**W**) for *gcs1* mutant at 3 DAP. (**X**) Higher magnification of (**W**). Although the main body of the ovule developed abnormally, the funiculus below the component still developed normally. Bars: 50 μm (**A**–**I**), 20 μm (**J**–**S**), 50 μm (**T**–**X**). CC: central cell, EC: egg cell, SC: synergid cell, DAP: days after pollination, SPC: sperm cell, PT: pollen tube.

Since part of the callose deposition area most likely overlapped with phloem unloading region (Karmann et al., 2018), it was hypothesized that callose deposition blocks the nutrition supply during seed formation. To test this hypothesis, cells of both WT and *gcs1* mutant ovules at 3 DAP were stained with aniline blue and observed using the *RPS5Apro::tdTomato-LTI6b* (Mizuta et al., 2015) membrane marker line. In WT, the signal was weak (Fig. 1T), and the ovule cells divided and expanded normally (Fig. 1U). However, in *gcs1* mutant, the signal remained intense (Fig. 1V), and the ovule cells were prevented from dividing and expanding (Fig. 1W). Although the main body of the ovule developed abnormally, part of the funiculus below the tissue still developed normally (Fig. 1X). These results suggest that a callose plug blocks substance transport to the end of the funiculus. However, after the fertilization, the block is removed and nutrients are delivered from the placenta to the ovule to form the seeds, functioning as a “gateway” to transport important substances from the parent body to the seed.

### Identification of callose-degrading enzyme genes encoding β-1,3-glucanase

Previous expression patterns (Kasahara et al., 2016) of callose-degrading enzyme genes were searched (Fig. 2A). The callose-degrading enzyme gene encodes β-1,3-glucanase, and is a part of a large gene family in *Arabidopsis*. A homology search based on the amino acid sequence of the typical gene, *β-1,3-glucanase 1*, revealed 49 genes in the *Arabidopsis* genome. Expression profiles of all genes were compared, and 47 transcripts were derived from expression of 36 genes detected during the first 48 h after fertilization in the WT. Of the 47 transcripts, 14 were found to be highly expressed at 48 h post-fertilization than before fertilization, and 11 transcripts were found to have peak expression at 48 h. Of these gene products, eight transcripts were not upregulated in *gcs1* mutant ovules, suggesting that these genes are associated with the deficiency of callose degradation in *gcs1* mutants (Fig. 2B). Among these eight, putative plasmodesmata-associated proteins (*AtBG_ppap*) (At5g42100) (Levy et al., 2007), *β-1,3-glucanase 1* (*BG1*) (At3g57270), and At4g29360 were annotated as the genes encoding callose degradation enzyme. Since the time-course fold-change of *AtBG_ppap* was the most significant, we further focused on this gene. Fig. S2A shows the structure of the *AtBG_ppap* gene. This gene consists of two exons, one of which is 1,209 bps, and the other is 69 bps, bridged by one intron (433 bps). CRISPR technology was applied to create an *Atbg_ppap* mutant that harbored a mutation site at 102 bps from ATG, resulting in a frameshift mutation (Fig. S2B). Figure S1C shows the amino acid (AA) sequence of the AtBG_ppap protein, containing 425 AAs. The *Atbg_ppap* mutant has an early termination of the AA synthesis, producing truncated protein of 87 AAs.

**Fig. 2.**
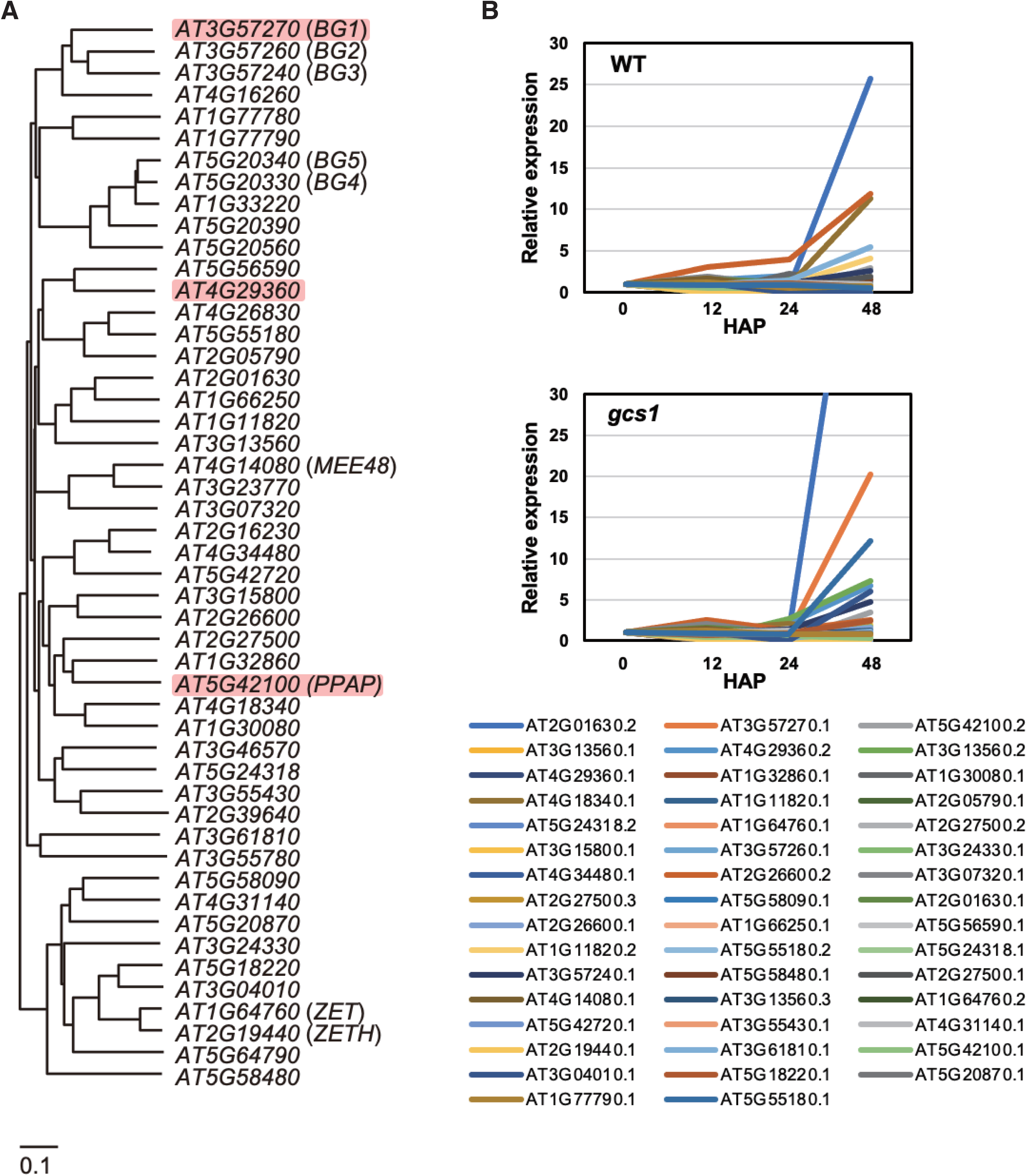
Phylogenetical analysis of ß-1,3-glucanase superfamily genes in *Arabidopsis*. **A**) Phylogeny of *ß-1,3-glucanase* gene family of *Arabidopsis.* A homology search was performed with tblastx using the CDS sequence of *ß-1,3-glucanase 1* (AT3G57270.1) as a query to obtain the transcripts with E values less than 10^-8^. Only one representative splice variant was employed for phylogenetic analysis. Multiple alignments were performed by Clustalw using the amino acid sequence of ß-1,3-glucanases, and a phylogenetic tree of 49 genes was generated by the neighbor-joining method, excluding outgroup genes. Genes with upregulated expression in WT and those repressed in *gcs1* mutant according to Fig. 1 are highlighted. (**B**) Expression of *ß-1,3-glucanase* genes during fertilization of WT and *gcs1* mutant plants. Transcriptomes were obtained by RNA-seq before (0) and 12, 24, and 48 h after pollination. Expression profiles were plotted for 47 transcripts of genes included in the phylogenetic analysis in (**A**), whose expression was detected at more than one timepoint, along with respective isoforms.

### Nutrition is blocked by callose deposition on the phloem end under unfertilized condition

Because callose deposition could block nutrient influx from the placenta to the ovule, subsequent experiments (Fig. 3) were designed to observe the influx of deposition blocked substance into the ovule using carboxy fluorescein diacetate (CFDA) (Oparka et al., 1994; Roberts et al., 1997). Live-cell imaging results showed that the carboxy fluorescein symplasmic tracer (CF) was transported to the ovule’s main body through the gate in WT ovules (Fig. 3A arrowheads, Video S2). However, it was not transported through the gate in *gcs1* mutant ovules (Fig. 3B arrowheads, Video S3), suggesting that nutrient flow from the funiculus was stopped by the gate in response to fertilization failure. In *Atbg_ppap* mutant ovules, the CF was barely observed in the main body (Fig. 3C arrowheads, Video S4), indicating that the nutrient flow was partially stopped by the gate, despite successful fertilization. Next, CF was observed in *AtBG_ppap* overexpression (*OEAtBG_ppap*) lines. Interestingly, the signal was detected in the main body, but was weaker than that observed in the WT and *Atbg_ppap* mutant at the part of the gate (Fig. 3D, Video S5), which may indicate that CF was more smoothly transported from the funiculus to the main body in the *OEAtBG_ppap* lines as the callose was constitutively degraded by AtBG_ppap. To investigate whether the *AtBG_ppap* gene is exclusively expressed in the ovule after fertilization, the expression patterns were examined by qRT-PCR from extracted various tissues. As shown in Fig. S3A and B, lower expression of *AtBG_ppap* was observed similarly both in the WT and *Atbg_ppap* (KO) lines at the WT (0DAP) and *gcs1* backgrounds (2DAP), however after fertilization (2DAP), *AtBG_ppap* was highly expressed in the WT but not in the KO line (Fig. S3C). As shown in Fig. S3D, *AtBG_ppap* was expressed in the cauline leaf, rosette leaf at low level, however in flower cluster and ovules (2DAP), *AtBG_ppap* was highly expressed. These results indicated that among tissues examined, the *AtBG_ppap* was highly expressed in the ovules at 2DAP.

**Fig. 3.**
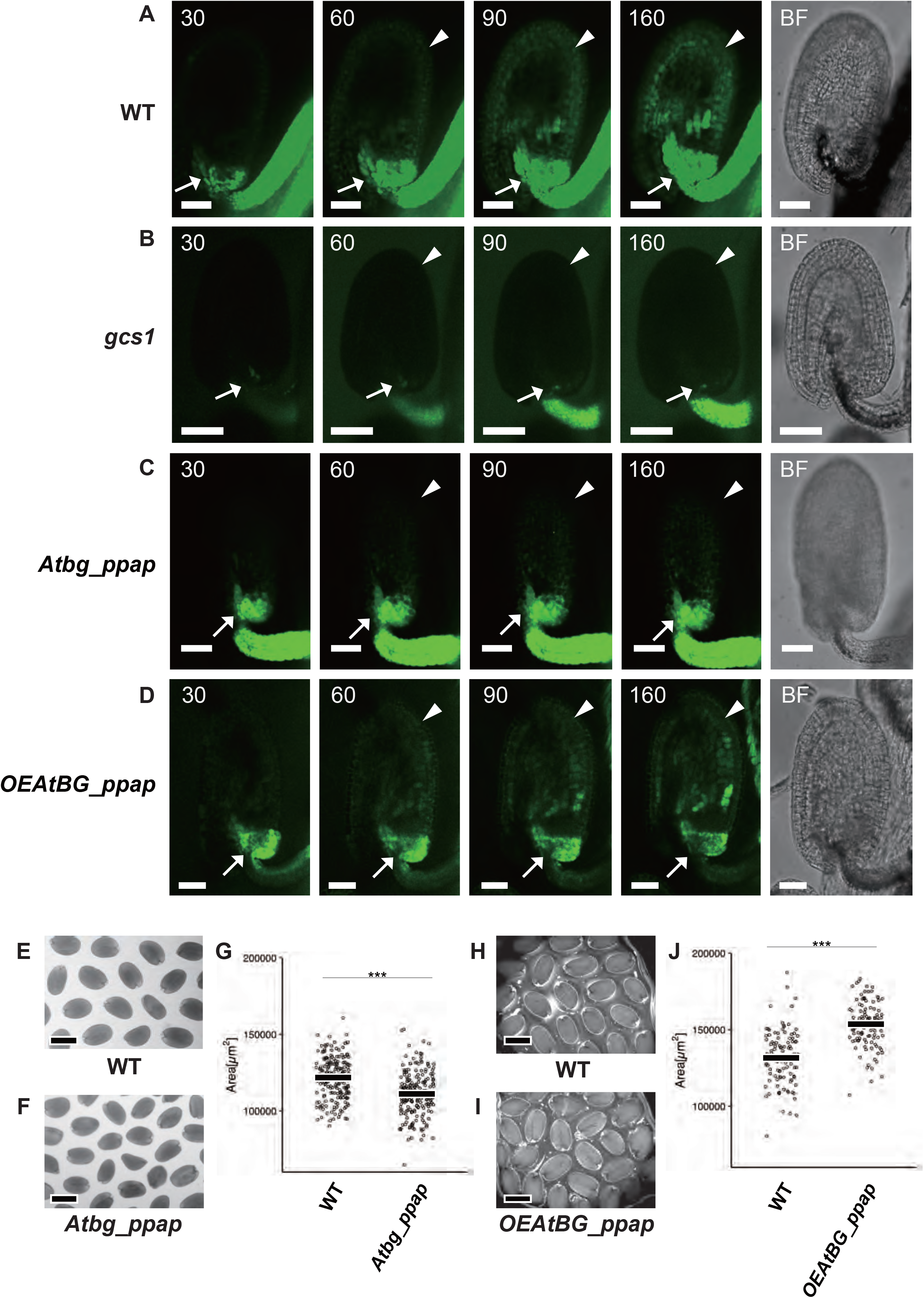
Nutrition flow is regulated by fertilization and AtBG_ppap, resulting in seed size modification. Live-cell imaging of CF dye tracer flow in the WT (A) at 30, 60, 90, and 160 min after absorption of CF (Video S2) and at bright-field (BF). CF dye tracer flow in gcs1 mutant (B) at 30, 60, 90, and 160 min after the absorption of CF (Video S3) and at BF. Live-cell imaging of CF dye tracer flow in *Atbg_ppap* mutant (C) at 30, 60, 90, and 160 min after the absorption of CF (Video S4) and at BF. Live-cell imaging of CF dye tracer flow in *OEAtBG_ppap* line (D) at 30, 60, 90, and 160 min after the absorption of CF (Video S5) and at BF. Arrows: Position of callose deposition. Arrowheads: Main body of the ovule. Actual seeds in the WT (E) and *Atbg_ppap* mutant (F). (G) WT seed size: 1.21 × 105 μm^2^ ± 0.13 × 105 μm^2^ (mean ± SD; n = 200 seeds), *Atbg_ppap* mutant seed size: 1.11 × 105 μm^2^ ± 1.45 μm^2^ (n = 200 seeds). *Atbg_ppap* mutant seeds were 8.4% smaller than those of WT. Welch’s two sample t-test was used to assess the significance of differences. ***P < 0.001. Actual seeds in WT (H) and *OEAtBG_ppap* (I). (J) WT seed size: 1.32 × 105 μm^2^ ± 0.18 × 105 μm^2^ (n = 200 seeds), *OEAtBG_ppap* seed size: 1.53 × 105 μm^2^ ± 0.15 × 105 μm^2^ (n = 200 seeds). *OEAtBG_ppap* seeds were 16.5% larger than those of WT. Welch’s two sample t-test was used to assess the significance of differences. ***P < 0.001. Scale bars: 50 μm (A–D) and 500 μm (E, F, H, I).

### *Atbg_ppap* produces smaller seeds and *OEAtBG_ppap* produces larger seeds

To further investigate the blocking effect, seed size of *Atbg_ppap* mutant was measured. Compared with the WT seeds (Fig. 3E), the seeds produced by the *Atbg_ppap* mutants were 8.4% smaller (Fig. 3F, G). Complementation test for the *Atbg_ppap* mutant was performed and showed that the seed size of the line was similar to that of WT (Fig. S3E). Next, to investigate whether the *Atbg_ppap* affected the other ovule phenotype, the ovule area and numbers were examined and resulted that those parameters had no significant differences between WT and *Atbg_ppap* lines (Fig. S3F and G). Similarly, the height of the plants had no significant differences (Fig. S3H). These results indicated that the *Atbg_ppap* mutation exclusively affected the seed size phenotype. In contrast, *OEAtBG**_**ppap* produced larger seeds than the WT, and the final seeds were 16.5% larger than those of the WT (Fig. 3H–J). To investigate whether the *OEAtBG**_**ppap* (OE) highly expressed in the ovules before and after fertilization, the expression patterns of WT and OE were compared by qRT-PCR (Fig. S4A and B). At both 0 DAP and 2DAP, OE expressed highly compared to WT. Next, to examine whether the OE line affects the ovule phenotype before fertilization, the size and numbers per one pistil were observed. At both these parameters, WT and OE had no significant differences (Fig. S4C and D). In addition, the plant height and sizes of one cotyledon, rosette leaf and cauline leaf were measured and among those parameters, there were no significant differences (Fig. S4E-H). Fig. S4I showed that OE seeds were larger than the WT seeds from 1DAP to 3DAP. These results indicated that the *OEAtBG**_**ppap* line exclusively affected the enlargement of the seed size.

These results strongly suggest that the callose degradation in *Atbg_ppap* mutant was incomplete and interfered with nutrient flow, leading to the development of smaller seeds compared to that of WT. Although callose blocked nutrient flow, mutant seeds were still formed, most likely because of the other redundantly functioning callose-degrading enzymes at the site. In contrast, in the *OEAtBG_ppap* seeds, callose was constitutively degraded, producing larger seeds because of the smooth nutrient transport, as shown in Fig. 3D and Video S5.

### Central cell fertilization is required to open the gate

Although our findings suggest that fertilization is required to degrade callose and open the gate for nutrients, the type of fertilization (egg or central cell) that initiates callose degradation is unclear. While, the *gcs1* mutant is useful for testing the morphological changes in the ovule during double fertilization failure, it is not applicable in the case of single fertilization, where one sperm cell fertilizes either the egg cell or central cell, but the other sperm cell does not. Hence, *kokopelli (kpl*) mutant pollen, which induces double fertilization at 28% ratio, egg cell single fertilization at 23% ratio, central cell single fertilization at 18% ratio, and no fertilization (POEM) at 24% ratio, was used for further assessment (Ron et al., 2010), since the *kpl* mutant is the only mutant that can induce single fertilization at present. A single fertilization event in either the egg or central cells was confirmed by aniline blue staining of each ovule at 2 DAP. In double-fertilized or central cell single-fertilized ovules, the callose was degraded (Fig. 4A, B). However, the signal was intense in egg cell single-fertilized ovules (Fig. 4C), suggesting that central cell fertilization is required for callose degradation. Since one of the prediction was that callose deposition could block the nutrient flow, the subsequent experiments were intended to demonstrate whether the deposition blocked substance influx to the ovule using CFDA (Oparka et al., 1994, and Roberts et al., 1997) after examining the backgrounds of *RPS5Apro::tdTomato-LTI6b* marker and *WOX2p::H2B-GFP + WOX2p::LTI-tdTomato* (Gooh et al., 2015) double markers for embryos, and the *AGL62::GFP* (Kang et al., 2008) marker for endosperm. An intense CF signal was observed above the phloem end of double-fertilized or central cell single-fertilized ovules (Fig. 4D–G or Fig. 4H–K arrowheads). However, this was not observed above the phloem end of the egg cell single-fertilized ovules (Fig. 4L–O), indicating that transportation did not occur if central cell fertilization failed (Fig. 4P). Conceptually identical results were obtained using esculin (Fig. S5), which indicates sucrose transport (Knox et al., 2018). An intense esculin signal was observed above the phloem end of double-fertilized or central cell single-fertilized ovules (Fig. S5A–C or Fig. S5D–F). However, this was not observed above the phloem end of the egg cell single-fertilized ovules (Fig. S5G–I) in comparison to WT ovules (Fig. S5J), nor was it observed in POEM ovules (Fig. S5K), indicating that sucrose transport did not occur when central cell fertilization of the ovule failed. These results show that sucrose and substance transport occurred only after the central cell fertilization (Fig. S5L), which was accompanied by callose degradation.

**Fig. 4.**
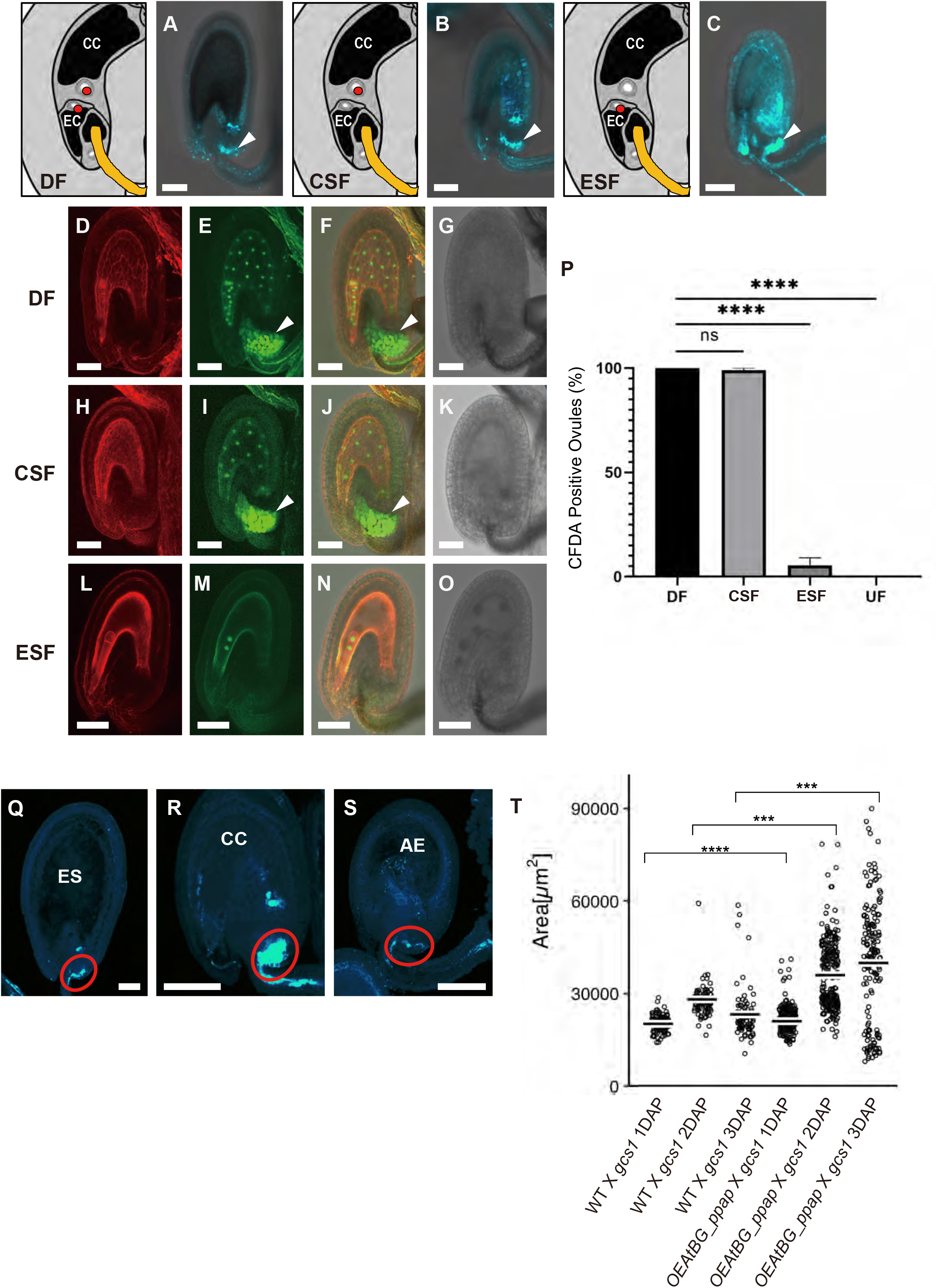
Central cell fertilization is required for callose degradation and nutrient flow. Aniline blue staining at 2 DAP (**A**), CC single fertilization (CSF) (**B**), and EC single fertilization (ESF) (**C**). CC: central cell; EC: egg cell. Arrowheads: Callose deposition. *WOX2p::H2B-GFP + WOX2p::LTI-tdTomat*o double marker (**D**), AGL62::GFP (**E**), merged (**F**), and bright-field (**G**) in DF ovules. *WOX2p::H2B-GFP + WOX2p::LTI-tdTomat*o double marker (**H**), AGL62::GFP (**I**), merged (**J**), and bright-field (**K**) in CSF ovules. *WOX2p::H2B-GFP + WOX2p::LTI-tdTomat*o double marker (**L**), AGL62::GFP (**M**), merged (**N**), and bright-field (**O**) in ESF ovules. Arrowheads: CF dye tracer signals. (**P**) Percentage of CF dye-tracer-positive ovules. DF: 100% ± 0% (n = 403 ovules). CSF: 98.5% ± 0.9% (n = 68 ovules). ESF: 5.8% ± 3.3% (n = 120 ovules). UF (Unfertilized): 0% ± 0% (n = 54 ovules). Dunnett’s multiple comparisons test was used to assess the significance of differences. ****P < 0.0001. ns = no significant. (**Q**–**S**) ES: endosperm, CC: central cell, AE: autonomous endosperm. Aniline blue staining results for WT (**Q**), *gcs1* mutant (**R**) and *fis2* mutant (**S**) ovules. Weak callose deposition in WT (**Q**), but intensive in *gcs1* mutant (**R**). (**S**) Weak callose deposition in *fis2* mutant without fertilization. (**T**) *OEAtBG_ppap* seeds without fertilization (POEM) were 7.2% (1 DAP), 30% (2 DAP), and 210% (3 DAP) larger than WT seeds without fertilization. Welch’s two sample t-test was used to assess the significance of differences. ***P < 0.001 and ****P < 0.0001. Bars: 50 μm (**A**–**O, Q**–**S**).

### *OEAtBGppap* produces larger seeds even without fertilization

An exceptional case of seed formation without fertilization is the apomixis (Koltunow and Grossniklaus, 2003; Underwood and Mercier, 2022). Mutant *fis2* is known to produce autonomous endosperm without fertilization (Luo et al., 1999). However, it is unclear why autonomous endosperm mutants produce seeds as large as those of the WT. Therefore, the relationship between the autonomous endosperm and the phloem end was further investigated using aniline blue staining. Callose was degraded in the WT but was deposited in *gcs1* mutant (Fig. 4Q, R); however, callose in *fis2* was degraded even without fertilization (Fig. 4S), indicating that autonomous endosperm formation was sufficient to remove callose and obtain nutrients via the phloem end. This result provides an answer to the question on why autonomous endosperm mutants produce seeds as large as those produced by the WT. To investigate whether *OEAtBG_ppap* (phloem end always open) can produce larger seeds, even without fertilization, the size of seeds from WT ovules crossed with *gcs1* mutant pollen and *OEAtBG_ppap* ovules crossed with *gcs1* mutant pollen were compared (Fig. 4T). Size of WT × *gcs1* mutant seeds was 2.11 × 10^4^ μm^2^ ± 0.28 × 10^4^ μm^2^ (n = 97 seeds) at 1 DAP, 2.89 × 10^4^ μm^2^ ± 0.46 × 10^4^ μm^2^ (n = 109 seeds) at 2 DAP, and 2.43 × 10^4^ μm^2^ ± 0.84 × 10^4^ μm^2^ (n = 82 seeds) at 3DAP. Size of *OEAtBG_ppap* × *gcs1* mutant seeds was 2.26 × 10^4^ μm^2^ ± 0.42 × 10^4^ μm^2^ (n = 240 seeds) at 1 DAP, 3.75 × 10^4^ μm^2^ ± 1.14 × 10^4^ μm^2^ (n = 249 seeds) at 2 DAP, and 5.11 × 10^4^ μm^2^ ± 3.41 × 10^4^ μm^2^ (n = 202 seeds) at 3 DAP. *OEAtBG_ppap* seeds without fertilization (POEM) were 7.2% (1 DAP), 30% (2 DAP), and 210% (3 DAP) larger than WT seeds without fertilization. These results indicated that *OEAtBG_ppap* may support further research on apomixis.

### Identification of the final form of the phloem end

We have identified the important function of the phloem end which is regulating seed nutrition by degrading callose after central cell fertilization in this study. However, the detailed phloem end structure is unclear, although its premature state has been described (Karmann et al., 2018). The final form of the phloem end was visualized using the phloem marker MtSEO2::GFP-ER (Noll et al., 2009; Froelich et al., 2011). At 1 DAE, the phloem end exhibited a blanched structure with two phloem tubes (Fig. 5A and Video S6), similar to the description by Karmann et al. (2018). However, at 2 DAP (equivalent to 3 DAE), the shape was further modified (Fig. 5B and Video S7), with a circular structure showing the final form of the phloem end in the plants. As the phloem end was identified and its position was likely to be identical to that of callose deposition, the spatial relationship between the two was further investigated.

**Fig. 5.**
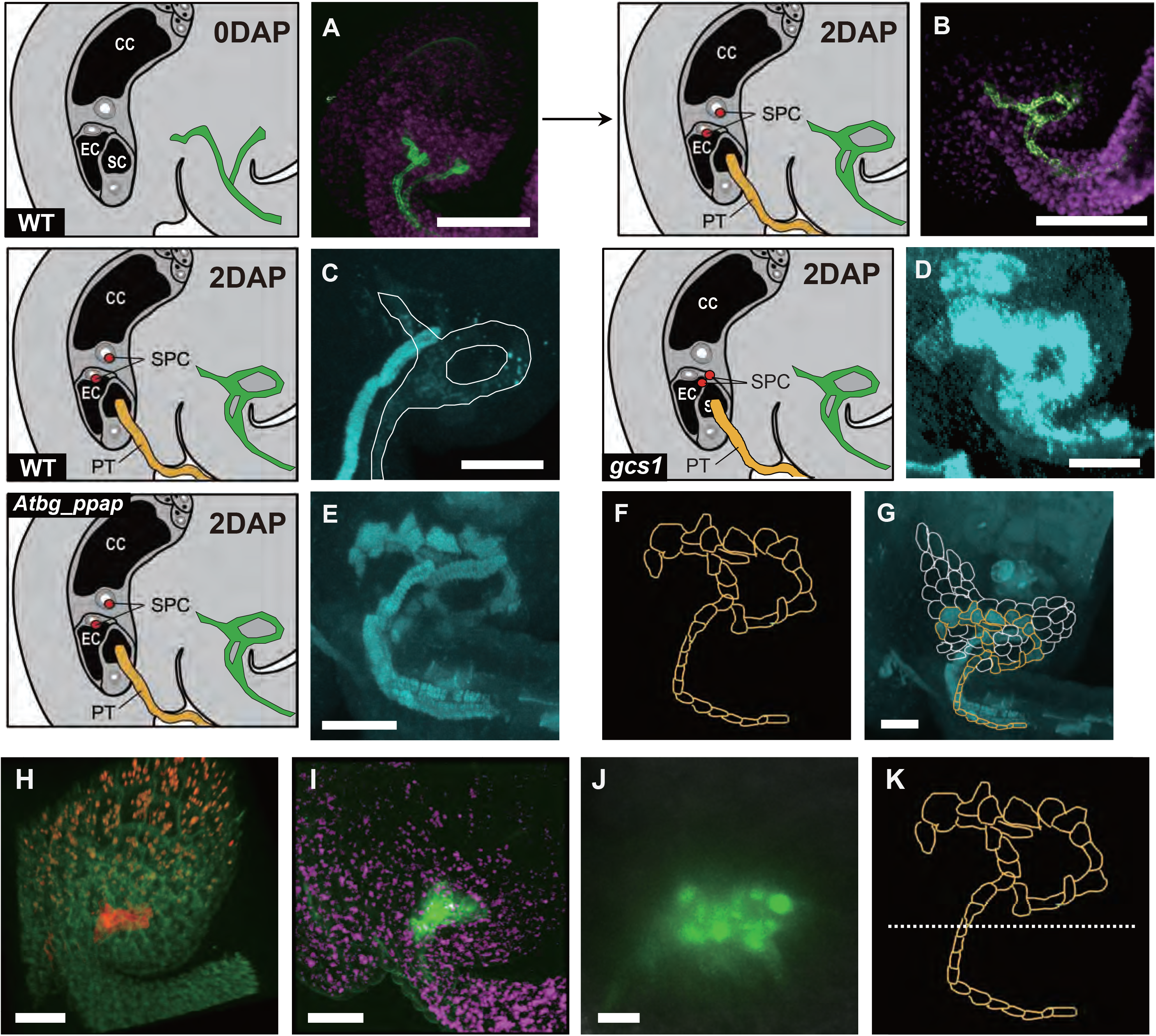
Detailed structure of the multi-functional phloem end of the ovule. (**A**) MtSEO2::GFP-ER (phloem marker) at 0 DAP indicating branched phloem tubes and (**B**) at 2 DAP indicating the final form of the phloem end of the ovule (PEO). Detailed structures of callose deposition parts in WT (**C**, outlined by white shape), *gcs1* (**D**), and *Atbg_ppap* (**E**) phloem end structure (**F**). (**G)** The structure with delineating peripheral cells depicted by white shape, designed from the Video S2 and S9. (**I**–**L**) Transmission electron microscopy images of ovule sections. The sieve element (SE) and companion cell (CO) near the chalaza of the ovules in the WT (**H**), *gcs1* mutant (**I**), *Atbg-ppap* mutant (**J**), and non-pollinated WT (NPT) (**K**). Arrowheads indicate plasmodesmata. The areas enclosed by the dashed rectangle are expanded in the insets. AtSWEET10-GFP was expressed at the phloem end **(L**–**N)** in WT at 2 DAP, exclusively in the upper half of PE (**O**). Bars: 50 μm (**A**–**E, G**), 1 μm (**H**–**K**; main panels), 0.2 µm (**H**–**K**; insets), 20 μm (**L, M**), and 10μm (**N**). CC: central cell, EC: egg cell, SC: synergid cell, DAP: days after pollination, SPC: sperm cell, PT: pollen tube.

The phenotype of the *Atbg_ppap* mutant (Fig. S2) was investigated using transcriptome data (Fig. 2) and aniline blue staining. Faint callose deposition, highlighted by the white line, was observed in the WT (Fig. 5C, Video S8), unlike the intense deposition in *gcs1* mutant (Fig. 5D). However, although the intensity of the callose deposition is moderate, the *Atbg_ppap* mutants showed persistent callose deposition, even after fertilization (Fig. 5E, Video S9). The structure of the callose deposit (Fig. 5E; sketched in Fig. 5F, Video S9) was identical to the final form of the phloem end (Fig. 5B, Video S7). Figure 5G illustrates the phloem end structure (yellow cells) and peripheral cells (white cells) of the phloem end from Fig. 3A, Video S2, and S9. These results suggest that symplasmic transportation from the funiculus to the ovule may be essential for the post-fertilization removal of callose deposit by AtBG_ppap at the phloem end.

Sucrose unloading in the ovule likely occurs through sucrose transporters. Previous studies (Kasahara et al., 2016) have shown the upregulation of SWEET sucrose transporters (Andrés et al., 2020) (Fig. S6) in the WT ovules after fertilization. Among the 17 *SWEET* genes in Arabidopsis, three genes, *SWEET1*, *7*, and *10*, were upregulated at 48 HAP in the WT, but downregulated or remained nearly flat at 24 to 48 h in *gcs1* mutant (Fig S6A). Phylogenetic analysis of these 17 *SWEET* genes showed that *SWEET1*, *7*, and *10* did not form a specific clade (Fig. S6B,C). Notably, in the WT, *SWEET10* promoter-driven GFP expression was observed exclusively at the gate (Fig. 5H–K, Video S10) and was limited to its upper part (Fig. 5K). Together with previous observations that phosphate and amino acid transporters (i.e., UmamiT; Müller et al., 2015) are also located at the gate in the ovule (Stadler et al., 2005; Werner et al., 2011; Müller et al., 2015; Vogiatzaki et al., 2017; Karmann et al, 2018), the phloem end identified in this report has at least two structural functions: site for nutrient unloading in the ovule and a gateway for callose deposition/degeneration. Figure 6 presents a summary of this report.

**Fig. 6.**
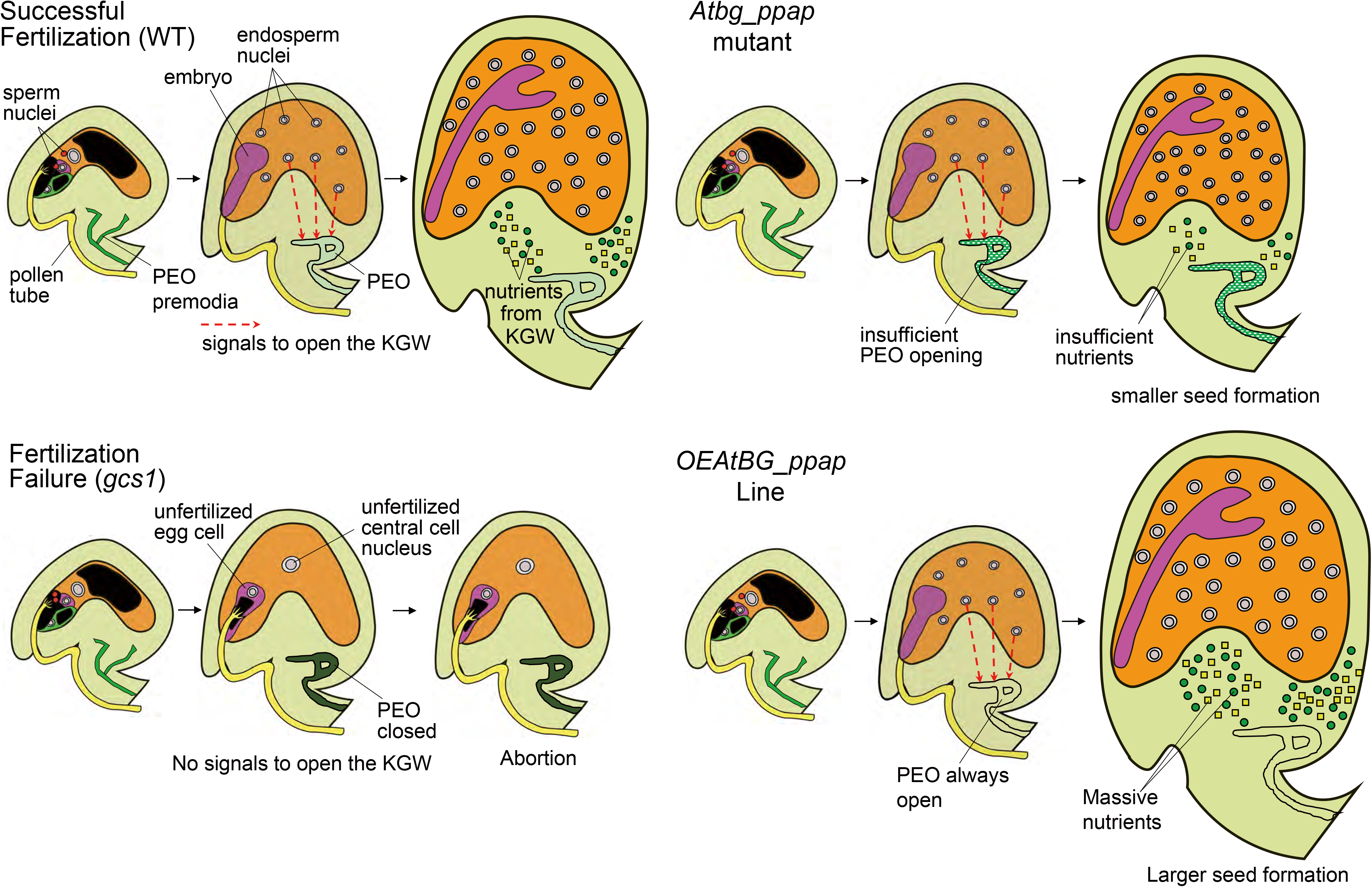
At 1 DAP, all ovule types (WT, *gcs1*, *Atbg_ppap*, and *OEAtBG_ppap*) contain phloem end (PE) primordia with branched phloem tubes; however mature PE is formed at approximately 2 DAP. In WT, after the central cell is fertilized, the cells (early endosperm) send fertilization completion signals to the PE. Subsequently, enzymes for callose degradation, including AtBG_ppap, degrade the callose plug in the PE, thus allowing the transport of nutrients to the fertilized ovule. In *gcs1* mutant, as the fertilization of the central cell fails, the unfertilized cell is unable to send fertilization completion signals to PE, and the callose degradation proteins are not synthesized; thus, the PE continues to deposit callose without transporting nutrients, which results in the death of the ovule. In *Atbg_ppap* mutant, the central cell is fertilized, and the early endosperm sends fertilization completion signals to the PE. However, as AtBG_ppap protein is not functional, callose degradation remains incomplete. The partial callose deposition interferes with the PE nutrient transport system, resulting in the formation of smaller seeds. In contrast, the *OEAtBG_ppap* line produces larger seeds than those produced by WT because *AtBG_ppap* is constitutively expressed and degrades callose in the PE, resulting in a flow of large amount of nutrients from the funiculus to the ovule.

## Discussion

### Identification of the new nutrition regulatory mechanism and AtBG_ppap

Seed size is one of the most important traits in plant breeding, because it is crucial for crop yield. In this study, we first identified *β-1,3-glucanase* genes through a homology search based on amino acid sequences. Expression profiles of all genes were compared and identified. *AtBG_ppap* gene among the tested genes was upregulated after fertilization in the WT but not in the unfertilized *gcs1* mutant ovules. Based on the results of the *Atbg_ppap* mutant and *OEAtBG_ppap* lines, we provide strong evidence for how and why AtBG_ppap modulates seed size. The gene encodes a membrane-bound protein involved in plasmodesmata gating *via* callose degradation (Levy et al., 2007). Our study showed that this gene is associated with seed size modulation. Our observations showed the deposition and degradation of callose before and after fertilization, respectively (Fig. 1), and indicated that callose deposition was distinguishable in NPT and *gcs1* mutant ovules, and these results led us to conclude that callose deposition continued until the ovule completed fertilization. The difference in the amount of callose deposition between NPT and *gcs1* mutant ovules could be explained by whether the ovule has pollen tube content (Kasahara et al., 2016). NPT ovules have no pollen tube contents because there are no pollen tubes, which means that there is no signal of failed fertilization. However, in the *gcs1* mutant ovules, because they have pollen tube contents, it is speculated that the ovule senses fertilization failure. The difference in the sensing mechanism between the NPT and *gcs1* mutant ovules most likely is responsible for the difference in the amount of callose deposition. However, the process by which NPT ovules accumulate callose without detecting the failure of fertilization needs further investigation.

In general, callose deposition is initiated at the chalazal end of an ovule at a very early stage of megaspore development (Rodkiewicz, 1970). Interestingly, an earlier observational study in *Hieracium* showed that callose deposition in sexual ovules was significantly higher than that in their apomictic counterparts (Tucker et al., 2001), and this is consistent with the conjecture derived from our study. However, their report also showed that, unlike sexual ovules, apomictic ovules have a higher callose accumulation at their chalazal ends than at their micropylar ends.

### Nutrients flow is governed by the phloem end and directly affects the seed size

In a previous study, Werner *et al*. (2011) had identified that before fertilization, the GFP protein cannot go beyond the phloem end of the ovule, but it can pass through it after fertilization. However, the reason why it could pass only after fertilization had not been investigated. We were able to investigate the precise nature of nutrient flow by using live-cell imaging of the CF dye tracer marker, as shown in Fig. 3. In *gcs1* mutant ovules, no signal was detected in the main body of the ovule, whereas *Atbg_ppap* mutant ovules showed a faint signal in the main body. These results suggest that in *gcs1* mutant ovules, the nutrient flow is completely blocked by the accumulated callose, resulting in seed abortion and no germination. However, in the *Atbg_ppap* mutant ovules, the nutrition is partially blocked by the callose deposition, resulting in production of smaller seeds with germination potential. The reason why the *gcs1* mutant ovules exhibit a complete shutdown of nutrient flow could be that the callose degradation enzyme is not upregulated in the absence of the fertilization event. In contrast, judging from the difference in the callose deposition amount between *Atbg_ppap* and *gcs1* mutant ovules, *Atbg_*ppap mutants can still produce smaller seeds. Since AtBG_ppap is a membrane-bound protein involved in callose degradation in the plasmodesmata (Levy et al., 2007), it is reasonable to assume that its defective mutants compromise the symplasmic system. This is in agreement with our observation of symplasmic tracer CFDA flow across the phloem unloading site in fertilized and unfertilized/failed to fertilize ovules. Furthermore, a similar observation made with esculin, a tracer for sugar transport, indicates that the initial nutrient (or at least sugar) flow to the ovule occurs mainly through the symplasmic pathway. However, both symplasmic and apoplasmic flows of nutrients have been proposed during fruit/seed development, with the latter initiating relatively later during the process (Lafon-Placette and Köhler, 2013; Ma et al., 2018). This could be the reason why the knock-out *Atbg_ppap* mutants still produced seeds, although with reduced size. Alternatively, unidentified redundantly acting AtBG_ppap-like proteins could play a role in this process. The presence of thick larger-than-cell-size callose deposition at the phloem unloading site in the NPT and *gcs1* mutant ovules observed in this study suggests that callose is actively deposited in the extracellular spaces. The differences between ovule/seed development in the NPT/*gcs1* and *Atbg_ppap* mutants also suggest that initial symplasmic flow is crucial for signal exchange between the reproductive and non-reproductive tissues, which could be linked to the initiation of symplasmic nutrient flow afterwards. Further research is required to confirm whether this is the case, and to understand the entire mechanism of nutrient flow during seed formation. From a commercial viewpoint, as *OEAtBG_ppap* ovules seemed to produce seeds of enlarged size, the gene could be useful for controlling seed size. Only one gene modification would be sufficient to increase and reduce the seed size in seed crops and fruit crops. Unlike, *AtBG_ppap*, the mechanism of action of most of the products of other seed size modulating genes, including plant hormone-related and cell metabolism-related proteins (Orozco-Arroyo et al., 2015, and Li et al., 2019) is not clearly understood yet. In contrast to these proteins, seed size modification by AtBG_ppap protein is completely explained by the nutrient flow governed by the callose blocking system. If a single amino acid transporter, UmamiT (Müller et al., 2015) is missing, 12% smaller seeds are observed. This observation is reasonable because these seeds lack a certain amount of amino acids. However, overexpression of UmamiT cannot be applied to enlarge seeds because seeds require various for nutrients, and only certain amounts of a single amino acid cannot compensate for all the nutrients. In contrast, overexpression of AtBG_ppap protein can be used to enlarge the seed because the protein physically removes callose deposit, which allows transportation of all nutrients from the phloem end to the seeds. So far, AtBG_ppap protein is the best candidate to modify the seed size by controlling the callose deposition/degradation.

### The blocking system avoids nutrient wastage

We previously discovered the fertilization recovery system that the second pollen tube compensates fertilization when the first pollen tube fails fertilization (Kasahara et al., 2012, Beale et al., 2012 and Kasahara et al., 2013). This is a wise strategy of plants to increase the number of seeds as a positive feedback system to increase the maximum usage of ovules by using the second pollen tube. In this study, we have identified another wise strategy of plants by using the callose-blocking system. In plants, supplying nutrients to unfertilized ovules can be quite wasteful because they cannot germinate, instead ensuring nutrients are supplied to fertilized ovules is important because this ensures that the next generation inherits the genetic information. To avoid nutrient wastage, plants appear to have evolved a system to block nutrient flow until the ovule senses the completion of fertilization as a negative feedback system. Our current study indicates that the initial symplasmic passage could be crucial for this process. Future research could shed more light on this aspect.

### Central cell fertilization is required to remove callose from the phloem end

This study showed that the fertilization completion signal originates from the fertilized central cell, but not from the fertilized egg cell (Fig. 4). Since egg cell fertilization produces an embryo that inherits genetic information, it is quite reasonable to expect it to send the fertilization completion signal. However, surprisingly, egg cell fertilization alone cannot induce opening of the gate. In contrast, central cell fertilization alone can trigger gate opening. Further, the reason behind endosperm cells sending the signal to the gate can be explained by their position in the ovule. After fertilization, it is the endosperm cell that is adjacent to the chalazal end, which is connected to the funiculus and callose accumulates between the funiculus and the endosperm, rather than the embryo cells, which are located on the micropyle side, the opposite side of the chalaza, which means that the energy cost to send the signal from the endosperm to the phloem end is lower than that from the embryo. Furthermore, unlike the endosperm, the early embryo consists of suspensor cells, often regarded as nutrition flow passages, at the micropylar end (Lafon-Placette and Köhler, 2014), which means that the energy cost to send nutrients from the suspensor cells is lower than that from the outside embryo sac. Moreover, the endosperm is essentially a tissue of nutrient reserve, which proliferates rapidly after fertilization, generating a greater ‘energy vacuum.’ This is another strong reason to speculate that endosperm cells generate signals to open the gate. It has long been discussed why autonomous endosperm mutants can produce larger ovules than embryo-only ovules. Polycomb group mutants, namely *mea* (Chaudhury et al., 1997, and Grossniklaus et al., 1998), *fis2* (Luo et al., 1999), *fie* (Ohad et al., 1999) and *msi1* (Guitton and Berger, 2005) mutants, can produce autonomous endosperms without fertilization. In this study, we have shown that in the *fis2* mutant, the autonomous endosperm exhibits less callose deposition at the chalazal end, resulting in larger ovule formation, most likely due to the signal flow from the proliferating endosperm cells. Another important finding of this study is that the levels of callose were similar in ovules with autonomous endosperm formation without fertilization and in ovules with central cell-only fertilization. Because the endosperm is the product of central cell fertilization, this correlation reinforces the link between callose degradation and central cell fertilization. We also observed that the *OEAtBG_ppap* ovule can be enlarged without fertilization by using *gcs1* mutant pollen. Similar to the case of the *fis2* mutant, the *OEAtBG_ppap* ovule can apparently degrade the callose without fertilization and obtain nutrients via the gate, thereby enlarging the seed size. This phenomenon could be applied to the enlargement of ovules without fertilization in various seed plants.

### Identification of the final form of the phloem end

Another important aspect of this study was the identification of the final structure of phloem tubes. Several previous studies have identified the phloem end as a loading region (Stadler et al., 2005; Werner et al., 2011; Müller et al., 2015; Vogiatzaki et al., 2017; Karmann et al, 2018), however, the structure of the final form of this region has not been properly identified. In this study, the region was identified as a site of callose deposition because the deposition site completely overlapped with the phloem end, indicating that this region not only functions as a nutrient unloading site but also regulates the nutrient supply by callose deposition/degradation. The starting point of the phloem translocation stream is inside the leaves of the plant body. The leaves synthesize sucrose by photosynthesis using CO_2_, H_2_O, and light energy from the sun and deliver it through the phloem to the entire plant. This study shows that the resources eventually arrive at the funiculus of the ovule and thereafter flow through the gate to be transported into the main body of the ovule. The gate is identified as the final form of the phloem end and its purpose is to regulate nutrient flow from the maternal placenta after receiving the fertilization completion signal. In future studies, the molecular mechanisms underlying the recognition of central cell fertilization and signaling that control callose deposition should be determined. Furthermore, the autonomous endosperm formation and gate control may be potential targets for apomixis induction in plants.

## Acknowledgments

We thank Maki Hattori, Aidi Xu, and Wei Xia for technical assistance. This work was supported by start-up funds from the School of Life Sciences, Fujian Agriculture and Forestry University (Grant #: 114-712018008 to R.D.K.) and the FAFU-UCR Joint Center, Haixia Institute of Science and Technology, Fujian Agriculture and Forestry University (Grant #: 102-118990010 to R.D.K.). This work was also supported by Chinese NSFC fund (Grant #: 31970809). This work was also supported by the Precursory Research for Embryonic Science and Technology (Grant #: 13416724 to R.D.K.; Kasahara Sakigake Project, Japan Science and Technology Agency). This work was partially supported by the Japan Society for the Promotion of Science Grants-in-Aid for Scientific Research (Grant #: 19H05361, 20H03273 and 21H00368 to M.Notaguchi, 16H06464 and 16K21727 to T.H.).

## Author contributions

M.Notaguchi, and R.D.K. conceptualized the study. X.L. and R.D.K. discovered the seed nutrition regulatory system. N.P.K. conducted all the live cell imaging experiments. X.L., P.B.A., M.Notaguchi, and R.D.K. designed the experiments. X.W., S.Z., and P.B.A. created *OEAtBG_ppap* and complementation lines. K.O. and R.D.K conducted the qRT-PCR experiments. T.I. created the *Atbg_ppap* mutant by CRISPR. X.L., N.P.K., X.W., S.Z., P.B.A., L.X., C.H., J.H., and R.D.K. performed the phenotypic analysis of *Atbg_ppap* mutant and *OEAtBG_ppap* line. M.Nakamura created the SWEET10 promoter-driven GFP line. K.K., and M.Notaguchi conducted the SWEET protein and the phloem end ultrastructure analysis. X.L., Y.S., and R.D.K. performed detailed phenotypic analysis for the phloem end callose deposition. X.L. and P.B.A. performed phenotypic analysis of esculin flow experiments. S.S., T.H., M. Notaguchi, and R.D.K. wrote the paper from the input of all authors’ experimental results.

## Materials and Methods

### Plant materials and growth conditions

*Arabidopsis thaliana* ecotypes Columbia (Col-0) was used as a WT plant. Testcross experiments were conducted in *gcs1*/*gcs1* (Mori et al., 2006; Nagahara et al., 2015), *RPS5Apro::tdTomato-LTI6b* (Mizuta et al., 2015), *kpl*/*kpl* (Ron et al., 2010), *WOX2p::H2B-GFP* + *WOX2p::LTI-tdTomato* double marker (Gooh et al., 2015), *AGL62::GFP* (Kang et al., 2008) and WT plants. Seeds were sterilized with 5% sodium hypochlorite containing 0.5% Triton X-100 and germinated on plates containing 0.5× Murashige and Skoog salts (pH 5.7) (Wako Pure Chemical), 2% sucrose, Gamborg’s B5 vitamin solution (Sigma), and 0.3% Gelrite (Wako Pure Chemical) in a growth chamber at 21.5℃ under 24 hours of light after cold treatment (4℃) for 2 days. Ten-day-old seedlings were transferred to Metro-Mix 350 soil (Sun Gro) and grown at 21.5℃ under 24 hours of light.

### Construction of *Atbg_ppap* mutant and *OEAtBG_ppap* lines

An *Atbg_ppap* mutant was made with CRISPR/Cas9 technology using an *Atbg_ppap*-targeting vector construct based on the pDe-Cas9 systems (Fauser et al., 2014). Gene-specific gRNA sequences unique for *AtBG_ppap* (At5g42100) was designed using tools available on the CRISPRdirect website (https://crispr.dbcls.jp/). gRNAs that contained restriction enzyme recognition sites around the Cas9 cleavage position were selected. Vector construction and generation of mutants were performed as described previously (Yamaguchi et al., 2017). The primer sequences used for this construction are shown in Table S1. Through selections with BASTA (Fujifilm Wako) and CAPS analyses, mutants containing 1 bp deletion (*Atbg_ppap-cr1*) in the open reading frame of *AtBG_ppap* was isolated. An *OEAtBG_ppap* line was made by following the method as described previously (Levy et al., 2007) by using the primer sets as shown in Table S1. For a rescue construct, the expression of *AtBG_ppap* was driven with 1571 bps of its native promoter and amplified from genomic DNA of Col ecotype background by using AtBG_PPAPpro_infu.fwd and AtBG_PPAPpro_infu.rev primer sets as shown in Table S1. Like in other constructs, GFP was sandwiched within the gene (after its functional domain and before the omega-site of the GPIanchor). The first part, fragment 1 followed by GFP (1158 bps) was amplified by using CVinfuPPAP.fwd and CVinfuPPAPpt1.rev primer sets (Table S1). The second part, fragment 2 after the GFP (599 bps) was amplified by using EGFP-AtBG_PPAPpt2_infu.fwd and EGFPAtBG_PPAPaltpt2_infu.rev primer sets (Table S1). The homozygous mutants of *Atbg_ppap*/*Atbg_ppap* were transformed with the Agrobacterium harboring the construct and phenotypic changes were observed in the transgenic lines.

### Construction of *AtSWEET10-GFP* line

The *AtSWEET10-GFP* construct was generated from a genomic clone of *AtSWEET10* (At5g50790), including a 2573-bp region 5’ upstream from the initiation ATG and a 1293-bp region 3’ downstream from the stop codon. GFP was fused to the *AtSWEET10* separated by a GGGGSGGGGSGGGS-linker (Shimozono and Miyawaki, 2008), just before the stop codon. The fluorescent protein-fusion construct was transferred to pBIN40 (Miyashima et al., 2011), which has been modified from pBIN19. Agrobacterium-mediated method (Clough and Bent, 1998) was used to generate transgenic plants. Primers used for this experiment are listed in Table S1.

### The phloem end phenotypic analyses

For staining of silique tissue, WT or other marker line flowers were emasculated at stage 12c (Christensen et al., 1997) and pollinated with WT, *gcs1*/*gcs1*, *kpl*/*kpl* or *Atbg_ppap*/*Atbg_ppap* pollen grains. Siliques were collected at 0, 1, 2, and 3DAP. After the samples were dissected and viewed with differential interference contrast microscopy, they were rinsed with Milli-Q (Millipore)–purified water and softened with 1 M NaOH for about 16 hours. The samples were directly stained with aniline blue solution [0.1% (w/v) aniline blue and 0.1 M K_3_PO_4_] for more than 3 hours. Confocal/two-photon images were acquired using a laser scanning inverted microscope (LSM780-DUO-NLO, Zeiss). The images were processed using the ZEN 2010 software (Zeiss) to create maximum-intensity projection images.

### CFDA and esculin

CFDA: A 100 μl drop of 5(6)-carboxyfluorescein diacetate (CFDA, optimum excitation 490 nm, emission 515 nm) (C8166, Sigma, stock solution 50 mg/ml in acetone) was applied to the cut edge of a single cotyledon at a concentration of 0.1 mg/ml in distilled water. Wrap the leaves in plastic wrap. The fluorescence of the dyes was visualized and recorded 4 hours after the application using a White Light Laser Confocal Microscope (Leica TCS SP8 X). WT, *Atbg_ppap*/*Atbg_ppap* or *OEAtBG_ppap*/*OEAtBG_ppap* line flowers were emasculated at stage 12c (Christensen et al., 1997) and pollinated with WT or *gcs1*/*gcs1* pollen grains. After 48 hours (=2DAP), the stem was cut with scissors at 5 cm from the root of the pollinated pistil. The cut plants were placed in PCR tubes containing 100 μl CFDA solution and incubated at 22℃ for 90 minutes. The ovary wall was cut off with a 27-gauge needle, and the stigma and placenta were split in two pieces. N5T medium was added dropwise to a glass bottom dish, ovules were lined up, and imaging was performed while sucking the solution. Time-lapse and Z-plane images were taken at 5-minute intervals, and 7 images were taken in 3 μm increments. Confocal images were acquired using an inverted fluorescence microscope (IX-83; Olympus) equipped with a disk scan confocal system (CSU-W1; Yokogawa Electric). Esculin: A 100 μl drop of 8 mg/ml esculin (66778-17-4, Sigma) (Esculin was dissolved in distilled water with 0.2% DMSO and 0.4% absolute ethyl alcohol) was applied to the cut edge of a single cotyledon. Wrap the leaves in plastic wrap. The fluorescence of the dyes was visualized and recorded 4 hours after the application using a White Light Laser Confocal Microscope (Leica TCS SP8 X).

## Supplemental information titles and legends

