## Supplementary material for "Seed size control via phloem end by callose deposition/degradation of β-1,3-glucanase": Fig. S2.pdf

**A**

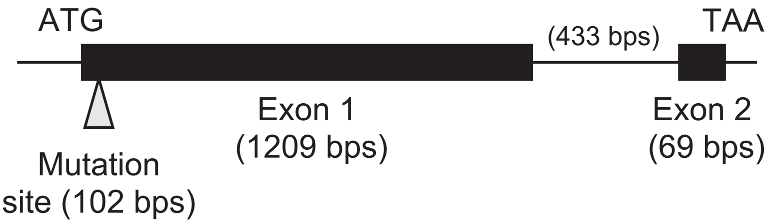


**B**

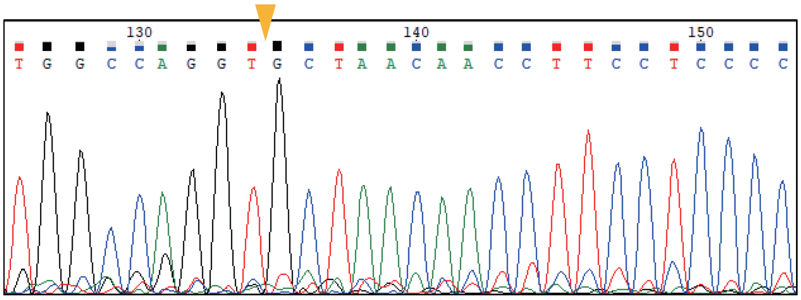


WT :TGGCCAGGTCGCTAACAACCTTCCTCCCC
Mutant :TGGCCAGGT--GCTAACAACCTTCCTCCCC

**C**

### AA sequence for WT

MASSSLQSLF SLFCLALFSL PLIVSSIGIN YGQVANNLPP PKNVIPLLKS VGATKVKLYD
ADPQALRAFA GSGFELTVAL GNEYLAQMSD PIKAQGWVKE NVQAYLPNTK IVAIVVGNEV
LTSNQSALTA ALFPAMQSIH GALVDCGLNK QIFVTTAHSL AILDVSYPPS ATSFRRDLLG
SLTPILDFHV KTGSPILINA YPFFAYEENP KHVSLDFVLF QPNQGFTDPG SNFHYDNMLF
AQVDAVYHAL DAVGISYKKV PIVVSETGWP SNGDPQEVGA TCDNARKYNG NLIKMMMSKK
MRTPIRPECD LTIFVFALFN ENMKPGPTSE RNYGLFNPDG TPVYSLGIKT SSTHSSGSGS
SNSTGGSSSG GGGNTGGSSS GGGIYQPVTG NPSPDYMSIS SAGGKGRFVE CVLFFFLLCI
IKLRL*

### AA sequence for *Atbg_ppap*

MASSSLQSLF SLFCLALFSL PLIVSSIGIN YGQVLTTFLP LKTSSLSSSL WELQKSSSMT
PIHKPYVPSP APASSSPWPS VTSTWLR*
