## Supplementary material for "Seed size control via phloem end by callose deposition/degradation of β-1,3-glucanase": Figure S legends.pdf

Figure S1. Callose signal intensities transition after fertilization and after failure of fertilization. In the cross of WT X WT, the callose intensity after 24 HAP, became weaker. In the cross of WT X *gcs1*, the intensity became stronger. Welch's two sample t-test was used to assess the significance of differences. ***P < 0.001.

Figure S2. Gene structure and a mutation site of a *Atbg_ppap* mutant.
(A) The *AtBG_ppap* (At5g42100) gene with the mutation site. Exon 1 contains 1209 base pairs (bps) and Exon 2 contains 69 bps. Intron between the Exon 1 and 2 contains 433 bps. The *Atbg_ppap* mutation contains 1 bp deletion at the position of 102th nucleotide from ATG. (B) Actual mutation site of the *Atbg_ppap* mutant. C was removed from the WT *AtBG_ppap* gene. (C) Truncated amino acid (AA) structure of *Atbg_ppap* mutant. WT *AtBG_ppap* protein contains 425 AAs. *Atbg_ppap* mutant produces 87 AAs.

Figure S3. *AtBG_ppap* expression in the ovules and the other tissues and phenotypic analysis in various tissues.
(A) The relative *AtBG_ppap* expression levels had no significant difference between WT and *Atbg_ppap* knockout (KO) lines. (B) The relative expression levels had no significant difference between WT and KO lines under no fertilization condition of the *gcs1* mutant background. (C) The *AtBG_ppap* was more highly expressed in the 2 DAP WT than in the KO background. Asterisks (* $P < 0.05$) indicate statistically significant differences relative to WT according to Student's *t* test. NS, not significant. (D) The relative *AtBG_ppap* expression levels among cauline leaf, rosette leaf, flower cluster and ovule at 2 DAP. No significant difference was detected between cauline leaf and rosette leaf but the expressions of flower cluster and ovule at 2 DAP were significantly higher than those of cauline leaf and rosette leaf. Asterisks indicate significant differences relative to WT (Student's t test, * p <0.05, ** p <0.01). NS; not significant. (E) Rescue construct (Comp Atbg) showed the similar seed size for the WT. Welch's two sample t-test was used to assess the significance of differences. ***P < 0.05. *Atbg_ppap* mutant showed the similar size in the ovule before fertilization (F), the similar number of the ovules per pistil (G) and the similar size in the plant height (H). Welch's two sample t-test was used to assess the significance of differences. NS; not significant.

Figure S4. *OEAtBG_ppap* plants produced larger seeds than those of WT.
Both (A) before fertilization (0DAP) and (B) after fertilization (2DAP), the *AtBG_ppap* expression in *OEAtBG_ppap* background (OE) was significantly higher than in WT background. Asterisks (* $P < 0.05$) indicate statistically significant differences relative to WT according to Student's *t* test. *OEAtBG_ppap* plants produced similar size to the ovules before fertilization (C), similar numbers of ovules before fertilization (D), similar sizes to plant height (E), one cotyledon (F), rosette leaf (G) and cauline leaf (H). Welch's two sample t-test was

used to assess the significance of differences. NS; not significant.

(I) WT 1DAP (WT ovule was crossed by WT pollen): $2.46 \times 10^4\ \mu m^2 \pm 0.6 \times 10^4\ \mu m^2$ (mean ± SD; n = 80 seeds), OEAtBG_ppap 1DAP (OEAtBG_ppap ovule was crossed by WT pollen): $2.63 \times 10^4\ \mu m^2 \pm 0.41 \times 10^4\ \mu m^2$ (n = 115 seeds). OEAtBG_ppap seeds were 6.9 % larger than WT seeds at 1DAP. WT 2DAP: $3.43 \times 10^4\ \mu m^2 \pm 0.5 \times 10^4\ \mu m^2$ (n = 93 seeds), OEAtBG_ppap 2DAP: $4.46 \times 10^4\ \mu m^2 \pm 0.77 \times 10^4\ \mu m^2$ (n = 152 seeds). OEAtBG_ppap seeds were 30 % larger than WT seeds at 2DAP. WT 3DAP: $6.14 \times 10^4\ \mu m^2 \pm 0.87 \times 10^4\ \mu m^2$ (n = 94 seeds), OEAtBG_ppap 3DAP: $6.93 \times 10^4\ \mu m^2 \pm 1.27 \times 10^4\ \mu m^2$ (n = 152 seeds). OEAtBG_ppap seeds were 12.9 % larger than WT seeds at 3DAP. Welch's two sample t-test was used to assess the significance of differences. *P < 0.05 and ***P < 0.001.

Figure S5. Direct sucrose unloading to the ovule main body by observing esculin flow.

DF: double fertilized, CSF: CC single-fertilized, ESF: EC single-fertilized. Esculin-flow was observed in DF (A−C), CSF (D−F), and ESF (G−I) ovules. AGL62::GFP (A), esculin (B), and bright-field (C) in DF ovule. AGL62::GFP (D), esculin (E) and bright field (F) in CSF ovule. AGL62::GFP (G), esculin (H), and bright-field (I) in ESF ovule. Esculin-flow in WT (J) and gcs1 (K). (L) Percentages of esculin-positive ovules. DF: 99.4% ± 0.3% (n = 335 ovules). CSF: 97.9% ± 2.1% (n = 94 ovules). ESF: 2.0% ± 2.8% (n = 99 ovules). UF (Unfertilized): 0% ± 0% (n = 34 ovules). Tukey's multiple comparisons test was used to assess the significance of differences. ****P < 0.0001. ns = no significant. Bars: 50 μm.

Figure S6. Expression of SWEETs during fertilization.

(A) Identification of SWEETs whose expression were upregulated in WT and repressed in gcs1 mutant plants during fertilization of Arabidopsis ovules. SWEET genes whose expression at 48 HAP were higher than those at 0, 12 and 24 HAP in WT, and increases from 24HAP to 48HAP were more moderate in gcs1 than in WT. (B, C) Phylogenetical analysis of SWEET family genes in Arabidopsis. (B) Phylogeny of Sugars Will Eventually be Exported Transporters (SWEETs) gene family of Arabidopsis. (C) Expression of SWEET genes during fertilization of WT and gcs1 mutant plants.
