## Supplementary figures and images for "Seed size control via phloem end by callose deposition/degradation of β-1,3-glucanase"

### Fig. S1.pdf

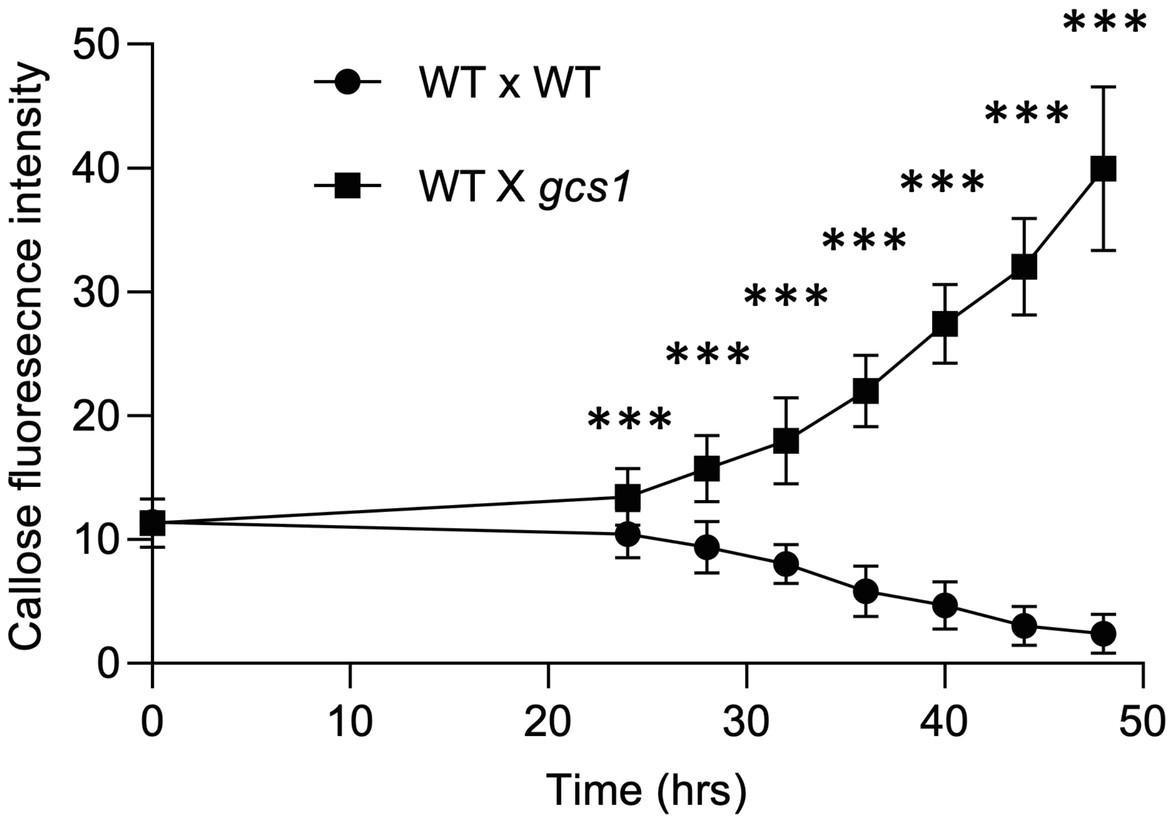

### Fig. S3.pdf

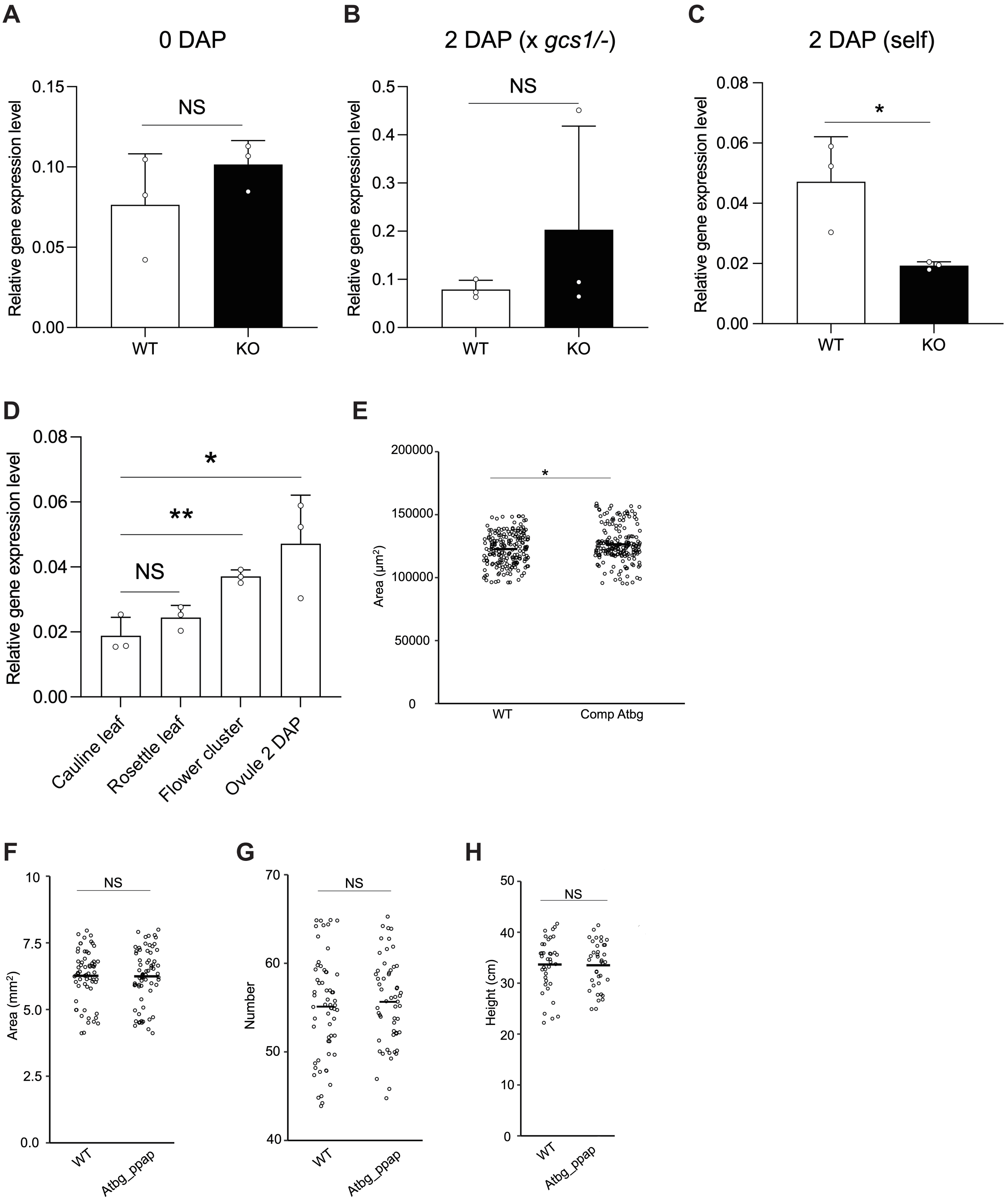

### Fig. S4.pdf

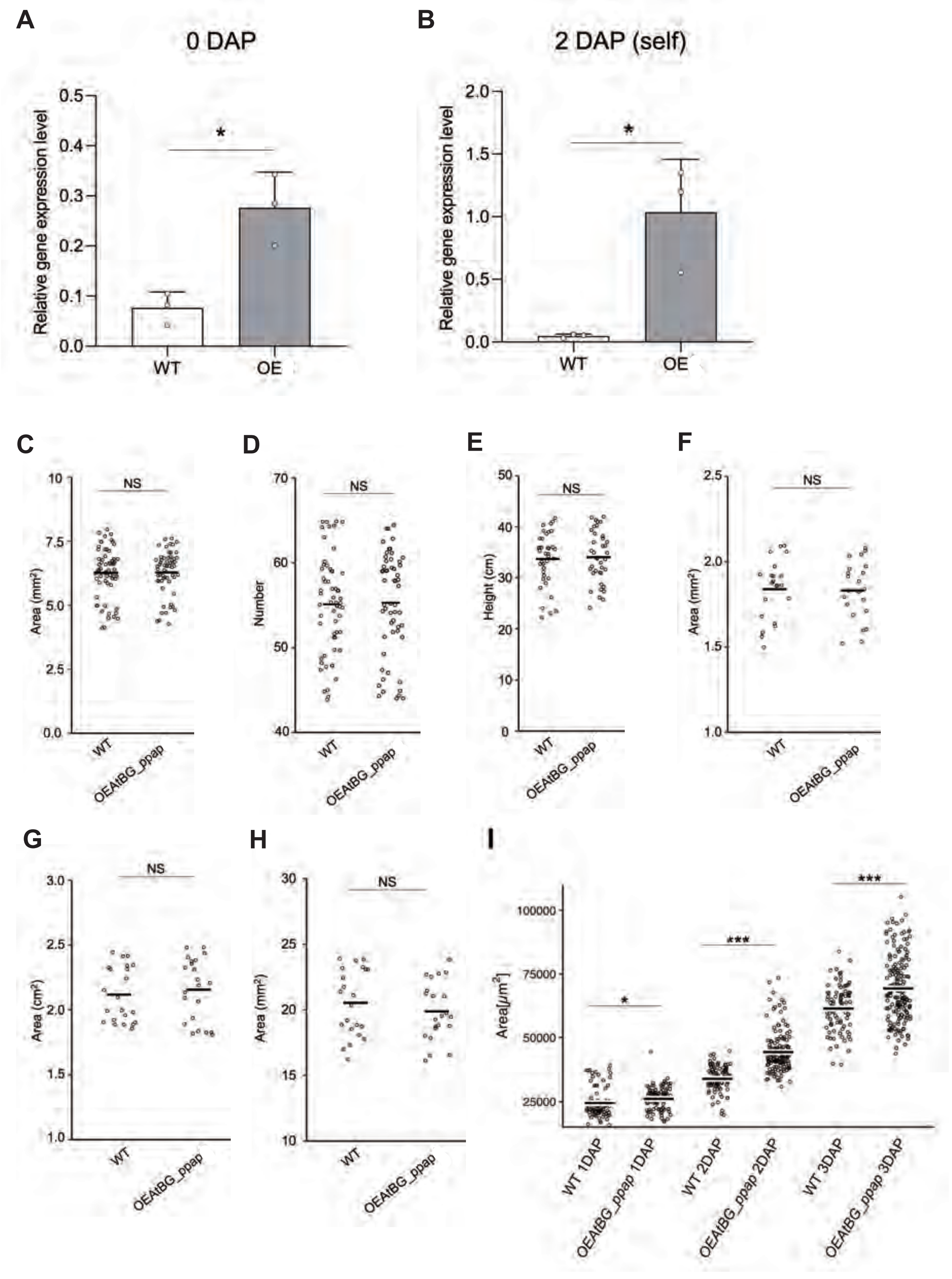

### Fig. S5.pdf

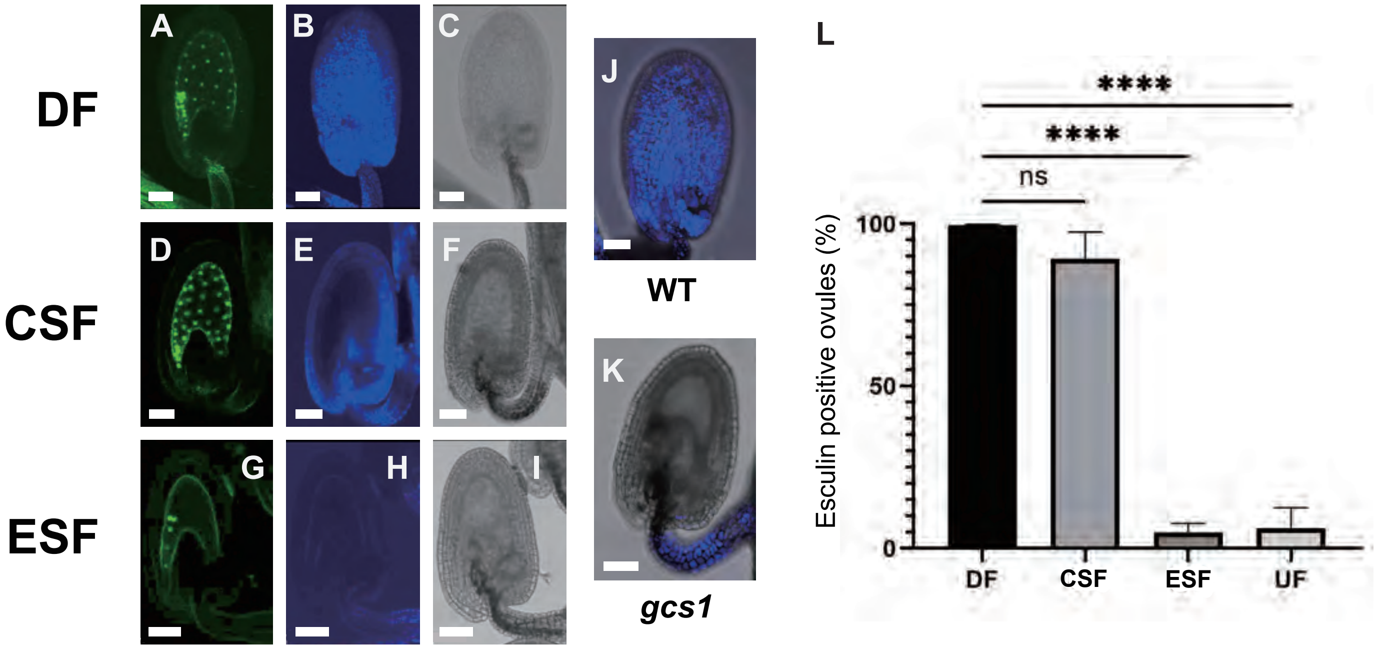

### Fig. S6.pdf

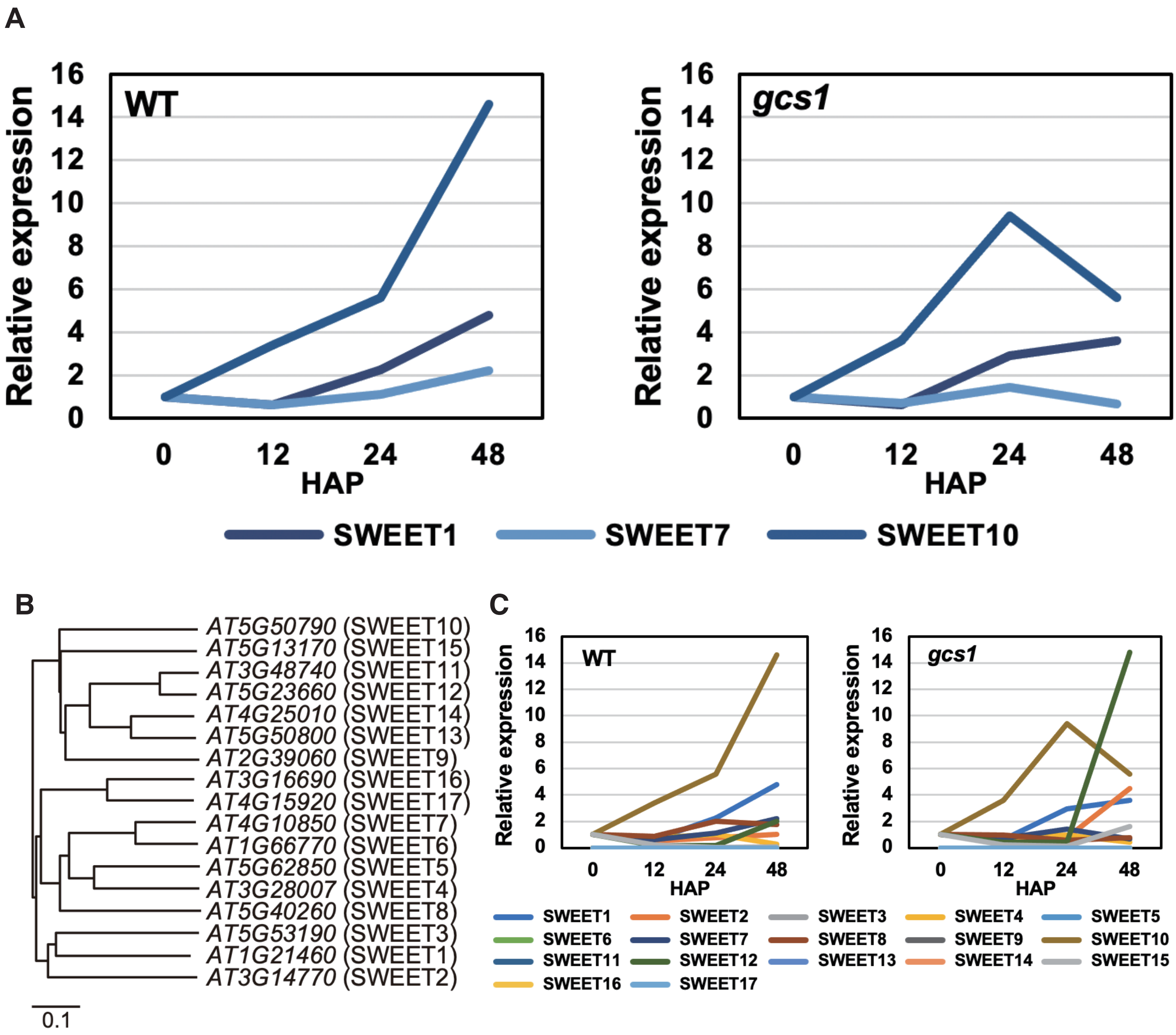
